# Rapid *in vitro* assays for screening neutralizing antibodies and antivirals against SARS-CoV-2

**DOI:** 10.1101/2020.07.22.216648

**Authors:** Jun-Gyu Park, Fatai S. Oladduni, Kevin Chiem, Chengjin Ye, Michael Pipenbrink, Thomas Moran, Mark R. Walter, James Kobie, Luis Martinez-Sobrido

## Abstract

Towards the end of 2019, a novel coronavirus (CoV) named severe acute respiratory syndrome coronavirus-2 (SARS-CoV-2), genetically similar to severe acute respiratory syndrome coronavirus-1 (SARS-CoV-1), emerged in Wuhan, Hubei province of China, and has been responsible of coronavirus disease 2019 (COVID-19) in humans. Since its first report, SARS-CoV-2 has resulted in a global pandemic, with over 10 million human infections and over 560,000 deaths reported worldwide at the end of June 2020. Currently, there are no United States (US) Food and Drug Administration (FDA)-approved vaccines and/or antivirals licensed against SARS-CoV-2, and the high economical and health impact of SARS-CoV-2 has placed global pressure on the scientific community to identify effective prophylactic and therapeutic treatments for the treatment of SARS-CoV-2 infection and associated COVID-19 disease. While some compounds have been already reported to reduce SARS-CoV-2 infection and a handful of monoclonal antibodies (mAbs) have been described that neutralize SARS-CoV-2, there is an urgent need for the development and standardization of assays which can be used in high through-put screening (HTS) settings to identify new antivirals and/or neutralizing mAbs against SARS-CoV-2. Here, we described a rapid, accurate and highly reproducible plaque reduction microneutralization (PRMNT) assay that can be quickly adapted for the identification and characterization of both neutralizing mAbs and antivirals against SARS-CoV-2. Importantly, our MNA is compatible with HTS settings to interrogate large and/or complex libraries of mAbs and/or antivirals to identify those with neutralizing and/or antiviral activity, respectively, against SARS-CoV-2.

## Introduction

A new variant of coronavirus (CoV) emerged around December of 2019 causing multiple hospitalizations in Wuhan, China (Bogoch et al., 2020; Choy et al., 2020). Ever since its first emergence, this novel severe acute respiratory syndrome coronavirus-2 (SARS-CoV-2) has been responsible for several cases of pneumonia leading to death among hospitalized patients all around the world. The transboundary spread of this coronavirus-associated acute respiratory disease, designated as coronavirus disease 2019 (COVID-19), prompted the World Health Organization (WHO) to declare the disease as a global pandemic on March 11^th^, 2020 (Organization). Besides the public health significance, COVID-19 has also disrupted the functioning of the global supply chain, bringing to a halt major economic activities all around the world.

SARS-CoV-2 is a close relative of severe acute respiratory syndrome coronavirus-1 (SARS-CoV-1) in the *Coronaviridae* family and has a single-stranded, non-segmented, positive-sense RNA genome (Schoeman and Fielding, 2019). Like other CoVs, SARS-CoV-2 encodes four conserved structural membrane (M), envelope (E), nucleoprotein (NP), and spike (S) proteins, all of which play an important role in viral replication (Masters, 2006; Mortola and Roy, 2004; Schoeman and Fielding, 2019; Wang et al., 2017). Of the four structural proteins, the S protein determines tissue tropism and host susceptibility to SARS-CoV-2 infection. An early aspect of SARS-CoV-2 replication in a susceptible host is the complex interaction between the viral S protein and the host cell surface receptor, the angiotensin converting enzyme 2 (ACE2). This intricate association enables viral fusion to the host cell membrane facilitating virus entry, an important requisite for successful virus replication. Owing to their importance in facilitating virus entry, the S protein has been a major target of neutralizing antibodies (NAbs) that are currently being produced against SARS-CoV-2.

Due to the high impact of COVID-19 and the lack of commercially available vaccines and/or NAbs, many United States (US) Food and Drug Administration (FDA)-approved or repurposed drugs are currently being suggested for the treatment of clinically-infected COVID-19 patients (Elfiky, 2020; Park et al., 2020b; Xu et al., 2020). As scientist and clinicians race to identify drugs for the treatment of SARS-CoV-2 infection and associated COVID-19 disease, some have been used in clinical settings with no supporting evidence of antiviral activity against SARS-CoV-2 (Reihani et al., 2020). The majority of these compounds are being recommended based on their ability to target specific steps of the replication cycle of similar viruses or, as seen in some cases, based on their ability to inhibit protozoan parasites (Yao et al., 2020). Apart from the lack of effective antiviral or immunotherapeutic activity against SARS-CoV-2, administration of certain antiviral compounds in COVID-19 patients has being a subject of safety concern among many clinicians. For instance, at high doses, chloroquine is extremely toxic causing death as a result of cardiac arrhythmias (Juurlink, 2020; Riou et al., 1988), and in the current pandemic, cases of chloroquine poisoning have been reported in Nigeria due to the abuse of the drug for treating COVID-19 patients (Chary et al., 2020). As a result, there is an urgent need for assays that can be used to screen the increasing lists of potential antiviral compounds and/or NAbs for an effective and safe anti-SARS-CoV-2 activity. Herein, we described a rapid approach that can be easily adapted for evaluating the potency and safety of a wide array of antiviral compounds and/or NAbs against SARS-CoV-2 in an *in vitro* setting.

## Materials

### Supplies and reagents

1. Vero E6 cells (BEI Resources, catalog # NR-596).
2. SARS-CoV-2 isolate USA-WA1/2020 (BEI Resources, catalog # NR-52281).
3. Dulbecco’s modified Eagle medium, DMEM (Corning, catalog # 15-013-CV).
4. Penicillin-Streptomycin L-glutamine 100x, PSG (Corning, catalog # 30-009-CI).
5. Fetal bovine serum, FBS (Avantor Seradigm, catalog # 1500-500).
6. Agar, 2% (Oxoid, catalog # LP0028).
7. Bovine serum albumin, 35% (BSA; Sigma-Aldrich, catalog # A7409).
8. BSA, 2.5% (Sigma-Aldrich, catalog # A9647).
9. DMEM/F-12 powder (Gibco, catalog # 12400-024).
10. Cell culture grade water (Corning, catalog # 25-055-CV).
11. Sodium bicarbonate, 5% (Sigma, catalog # S-5761).
12. DEAE-Dextran, 1% (MP Biomedicals, catalog # 195133).
13. Crystal violet, 1% (Fisher Scientific, catalog # C581-100).
14. Avicel PH-101, 1% (Sigma-Aldrich, catalog # 11365).
15. Formalin solution, neutral buffered, 10% (Sigma-Aldrich, catalog # HT501128-4L).
16. Triton X-100, 0.5% (Sigma-Aldrich, catalog # X100-500ML).
17. Mouse anti-SARS-1 nucleoprotein (NP) mAb 1C7 generated at the Center for Therapeutic Antibody Development at The Icahn School of Medicine at Mount Sinai (ISMMS) (Millipore Sigma, catalog # ZMS1075).
18. Remdesivir (AOBIOUS, catalog # AOB36496).
19. Human monoclonal antibody (hmAb) 1207B4 produced in house.
20. VECTASTAIN^®^ ABC-HRP Kit, Peroxidase (POD) (Mouse IgG) (Vector Laboratory, catalog # P-4002).
21. DAB Substrate Kit, POD (HRP), with Nickel (Vector Laboratory, catalog # SK-4100).
22. IRDye 800CW goat anti-mouse IgG secondary antibody (LI-COR, catalog # 926-32210).
23. DRAQ5^™^ Fluorescent Probe Solution 5 mM (Thermo scientific, catalog # 62251).
24. 6-well cell culture plate (Greiner Bio-One, catalog # 657160).
25. 96-well cell culture plate (Greiner Bio-One, catalog # 655180).
26. Polystyrene tissue culture flask (Corning, catalog # 431081).
27. Polypropylene sterile conical tube, 15 mL (Greiner Bio-One, catalog # 188261).
28. Polypropylene sterile conical tube, 50 mL (Greiner Bio-One, catalog # 227270).
29. Serological pipette, 5 mL (Greiner Bio-One, catalog # 606 180).
30. Serological pipette,10 mL (Greiner Bio-One, catalog # 607 180).
31. Serological pipette, 25 mL (Greiner Bio-One, catalog # 760 180).
32. Universal pipette tip, 20 μL (VWR, catalog # 76322-134).
33. Universal pipette tip, 200 μL (VWR, catalog # 76322-150).
34. Universal pipette tip, 1000 μL (VWR, catalog # 16466-008).
35. Microcentrifuge tube, 1.5 mL (VWR, catalog # 89000-028).
36. Sterile basin (VWR, catalog # 89094-680).
37. CO_2_ incubator (PHCbi, model # MCO-170AICUVDL).
38. CTL ImmunoSpot plate reader and counting software (Cellular Technology Limited).
39. Odyssey Sa Infrared Imaging System (LI-COR, model # 9260).
40. GraphPad Prism (GraphPad Software Inc., version 8.0).

### Media type and recipes

41. Cell-maintenance media (DMEM supplemented with 10% FBS and 1% PSG).
42. Post-infection media (DMEM supplemented with 2% FBS and 1% PSG).
43. Infection media (DMEM supplemented with 1% PSG).
44. DMEM/F-12/Agar mixture (DMEM-F12 with 1% DEAE-Dextran, 2% agar, and 5% NaHCO_3_).
45. Plaque reduction microneutralization (PRMNT) media (Post-infection media with 1% Avicel).

#### Biosafety Recommendations

Although SARS-CoV-2 has been temporarily classified as a category B pathogen, biosafety caution applicable to the category A pathogens are strongly recommended when handling this pathogen. Manipulation of SARS-CoV-2 should only be carried out in a biosafety level 3 (BSL3) facility with an appropriate engineering system designed to produce negative air pressure and enhance the safety of laboratory workers. All individuals working with SARS-CoV-2 must have undergone proper biosafety training to work at BSL3 laboratories and be cleared proficient to carry out procedures requiring working with the virus. Cell culture procedures are carried out in BSL2 and moved to BSL3 when ready for viral infection.

#### Cells

Vero E6 cells, which have been demonstrated to be permissive to SARS-CoV-2, is the preferred cell type for the assays described in this protocol (Imai et al., 2020). Other cells types such as Vero E6 transfected to express transmembrane protease, serine 2 (TMPRSS2), a serine protease which activates SARS-CoV-2 infection (Matsuyama et al., 2020), can also be used. An important consideration when choosing a cell line for this assay is the ability of SARS-CoV-2 to efficiently infect and replicate in the cell line. It is important to seed cells a day before the assay to achieve approximately ~85-95% confluency. Vero E6 cells are maintained in DMEM supplemented with 10% FBS and 1% PSG. We recommend using a low passage of Vero E6 cells as we have found that these cells grow better and are more viable at low passage. It is equally important to make sure the cells are free from any contaminants including mycoplasma.

#### Virus

For the experiments described in this manuscript we used the SARS-CoV-2 isolate USA-WA1/2020 available from BEI Resources. For the proposed experiments, we recommend using ~100-200 plaque forming units (PFU)/well for a 96-well plate. Higher concentrations of the virus may affect the sensitivity of the assay while using lower viral concentrations may result in reduced number of plaques/well. Following the experimental conditions described in this manuscript, you would expect to have ~100-200 positive staining cells/well at 24 h p.i.

#### Serum samples

It is important to heat-inactivate serum or plasma samples from COVID-19 patients or SARS-CoV-2-infected animals at 56°C for 1 h before performing the PRMNT assay to destroy complement proteins or residual SARS-CoV-2 particles. The experimental procedures to inactivate the virus should be confirmed and approved by the institutional biosafety committee (IBC) of the institute. It has been reported that complement deposition on virus envelope may lead to infection-enhancement which may mask the neutralizing effects of Abs contained in serum or plasma samples (Montefiori, 1997). For serum samples or plasma samples, we recommend starting with a 1:100 dilution to avoid potential impurities that may affect the sensitivity of the assay. In the case of mAbs, we recommend staring with 10 μg. However, this will depend on the neutralizing capability of the mAb. We used a hmAb, 1207B4, for the purposes of this study.

#### Drugs

The drug used in this study is remdesivir, a nucleoside analog which interferes with the replication of viral genome by inhibiting viral RNA-dependent RNA polymerase (RdRp) (Gordon et al., 2020). Any other antiviral drug or compound with potential antiviral activity against SARS-CoV-2 can equally be used in the assays described below.

#### Controls

It is important to validate the described PRMNT assays by including proper positive and negative controls. A mAb with a known neutralizing titer or serum and/or plasma samples from convalescent SARS-CoV-2 patients that contains known NAbs can be used as positive controls. In the case of the antiviral drugs, we recommend to use remdesivir since it has been shown to have antiviral activity against SARS-CoV-2 (Choy et al., 2020; Ferner and Aronson, 2020). Negative controls include the use of SARS-CoV-2-infected Vero E6 cells without NAbs, sera, or drugs. An additional recommended control to validate the immunostaining with the NP 1C7 mAb is to include cells without viral infection.

#### General Procedure

All cell cultures procedures and antibody/antiviral dilutions are generally performed in the BSL2 before transfer to the BSL3 facility on the day of virus infection. For PRMNT assay, the amount of virus per well can be optimized based on the virus titer in the stock. During the inactivation procedure, it is important to ensure that the entire plate and its lid are fully submerged in 10% formalin solution for proper inactivation. After the inactivation procedure plates can be transferred from BSL3 to the BSL2 for immunostaining and development.

## Results

### Preparation of SARS-CoV-2 stock

1. A day before viral infection, seed Vero E6 cells in a T-225 cell culture flask in cell-maintenance media to attain ~85%-95% confluency on the day of infection and place them in a humidified incubator at 37°C with 5% CO_2_.
2. Thaw the stock virus and prepare an infection media containing the virus inoculum at a MOI of 0.001.
3. Remove cell maintenance media and replace with infection media containing the virus in a total volume of 10 mL. Place the tissue culture flasks of infected cells in Ziploc bags and transport them on a flat tray to the humidified incubator and incubate at 37°C in the presence of 5% CO_2_. Note: shake the flasks, gently, every 10 minutes to prevent cells from drying out during a 1 h incubation time-point (TP).
4. After 1 h of virus adsorption, in a humidified incubator at 37°C with 5% CO_2_, remove the virus inoculum and replace with ~40 mL of post-infection media.
5. Incubate for 72 h in a humidified incubator at 37°C with 5% CO_2_. Note: SARS-CoV-2 behaves differently compared to most other viruses in that the presence of cytopathic effect (CPE) does not correlate with increased virus titer. For our virus stock, we carried out a multistep growth kinetics to determine the best time with the highest virus yield. In our hands, this correlated nicely with low CPE at ~72 h of incubation.
6. Freeze-thaw the flask containing the tissue culture supernatant (TCS) once, followed by centrifugation at 2,000 g for 10 min at 4°C.
7. Aliquot the TCS in 500 μL volumes, and store at - 80°C until further use.

### Determining the infectious virus titer by plaque assay

1. The day before infection, prepare Vero E6 cells (6-well plate format, 1 x 10^6^ cells /well) in cell-maintenance media and place them in a humidified incubator at 37°C with 5% CO_2_.
2. Prepare 10-fold serial dilutions of the SARS-CoV-2 stock in infection media. Briefly, pipette 450 μL of infection media and add 50 μL of the virus stock in the first tube. Transfer 50 μL from tube #1 (10^-1^ dilution) to tube #2 (10^-2^ dilution) and continue doing serial dilutions. Note: It is important to change the tips between viral dilutions to prevent the transfer of virus particles from a lower to higher diluted infection medium.
3. Infect the 6-well plates of Vero E6 cells with 200 μL of each of the SARS-CoV-2 dilutions and incubate for 1 h in a humidified incubator at 37°C with 5% CO_2_. Place the plates of infected cells in a Ziploc bag and transport them on a flat tray to the humidified incubator and incubate at 37°C in the presence of 5% CO_2_.
4. To prevent cells from drying and to also facilitate virus adsorption, shake the plates gently every 10 minutes.
5. After 1 h of viral absorption, remove the virus inoculum and overlay the cells with DMEM/F-12/Agar mixture. Note: ensure that at the start of the infection, the 2% agar is melted in a microwave and the temperature brought down to approximately 42°C when mixing with warm DMEM/F-12/Agar mixture. Note: It is important to keep the agar cool enough not to burn the cells, and at the same time warm enough not to solidify in the process of pipetting.
6. Allow the agar to solidify and invert the plates to prevent the accumulation of moisture during incubation.
7. Incubate the cells for 72 h in a humidified incubator at 37°C in the presence of 5% CO_2_.
8. At 72 h post-infection (h p.i.), fix infected cells in 10% formalin solution for 24 h. Note: After 24 h fixation with 10% formalin solution, plates can be moved from the BSL3 to the BSL2 to complete the viral titration.
9. Perform immunostaining using the cross-reactive SARS-CoV-1 NP mAb 1C7, at a working concentration of 1 μg/mL. Note: Alternatively, cells can also be stained with 1% crystal violet for plaque visualization. Count the number of plaques, and calculate the virus titer in plaque forming units per mL (PFU/mL).

### A plaque reduction microneutralization (PRMNT) assay to identify Abs with neutralizing activity against SARS-CoV-2

This assay can be used to evaluate SARS-CoV-2 neutralizing activity by mAbs, polyclonal antibodies (pAbs) or by an Ab-containing sample such as serum plasma from mammalian host species. Since serial dilutions of the Ab containing samples are used, the PRMNT assay measures the neutralizing activity of Abs in a concentration dependent manner. The ability of a NAbs to inhibit virus infection is manifested in the reduced capacity of the virus to produce visible plaques when compared to virus-only infected control cells. The PRMNT assay is similar to a plaque reduction neutralization (PRNT) assay with the only difference that the PRMNT utilizes 96-well plates. Although in this manuscript we describe the use of PRMNT in 96-well plates, similar experimental approaches can be miniaturized and conducted in 384-well cell culture plates for HTS purposes.

### A PRMNT assay to identify NAbs

The process of entry into a susceptible host cell is an important determinant of infectivity and pathogenesis of viruses, including CoVs (Li, 2016; Perlman and Netland, 2009). SARS-CoV-2 relies on the ability of its S glycoprotein to bind to the ACE2 receptor through its receptor-binding domain (RBD) driving a conformational change that culminates in the fusion of the viral envelope with the host cell membrane, and cell entry (Shang et al., 2020). SARS-CoV-2 S is made up of 2 subunits, S1 and S2. During pre-treatment, mAbs, or Ab containing samples, are pre-incubated with SARS-CoV-2 at 37°C for 1h. This enables the Abs to bind to the S protein blocking virus attachment, and subsequently interfering with the process of virus entry. While some SARS-CoV-2-induced NAbs can preferentially bind to either the S1 or S2 subunit of the S protein, others can bind, with a high affinity, to the RBD located within the S1 subunit (Wang et al., 2020; Wu et al., 2020). After 1 h of incubation, the Ab-virus mixture is then transferred to a confluent (~85-95%) monolayer of Vero E6 cells and incubated for 24 h at 37°C. During this period, the unbound virus particles are able to attach to cells and initiate virus replication which can be determined by staining for SARS-CoV-2. In the post-treatment condition, virus adsorption is allowed to progress for 1h at 37°C. This gives an opportunity for the virus to initiate viral entry by binding to the cell surface receptor. NAb incubated with the infected cells may interfere with active viral replication partly by blocking later steps of the virus entry into the cell or by inhibiting the cell-to-cell spread of virus progeny. In both pre- and post-treatment conditions, the extent of virus neutralization can be visualized upon staining with a cross-reactive anti-SARS-CoV-1 NP mAb (1C7), and the neutralization titer 50 (NT50) is calculated using a sigmoidal dose-response curve. The NT_50_ of an Ab is the dilution which inhibited viral replication in 50% of the infected cells. While the pre-treatment PRMNT assay can be described, to some extent, as a prophylactic assessment of a NAb, the post-treatment PRMNT assay offers a therapeutic evaluation of the neutralizing activity of Abs against SARS-CoV-2. In addition, while the pre-treatment PRMNT assay focuses on identifying NAbs targeting the SARS-CoV-2 S RBD, the post-treatment PRMNT assay also allow the identification of Abs targeting other regions on the viral S glycoprotein involved in the fusion event (e.g. S2), or Abs affecting other steps in the replication cycle of the virus (e.g. virus assembly and/or budding). In general, a potent SARS-CoV-2 NAb will show a low NT_50_ titer whereas a weak NAb will give a high NT_50_ value. This protocol precludes the necessity for a cytotoxicity assay since Abs are not known to produce cytotoxic effects in cell cultures. However, a potential toxic effect of the NAbs can be determined when developing the assay with the infrared staining described below.

### Pre-treatment PRMNT assay

1. Seed ~1 × 10^4^ Vero E6 cells/well the day before virus infection in 96-well plates using cell maintenance media.
2. On the day of infection, check the confluency of the Vero E6 cells under a light microscope. The optimal cell confluency should be ~85-95%.
3. Prepare a 2-fold serial dilution of the Ab (or Ab containing sample) in an empty, sterile 96-well plate using infection media. Briefly, add 50 μL of infection media to columns 2 to 12, and add 100 μL of the desired starting concentration of each Ab, or Ab containing sample, to column 1. Transfer 50 μL of Ab from column 1 to column 2, and mix ~10 times using a multi-channel pipette. Repeat this process from column 2 to column 10, changing the tips between dilutions to prevent transfer of residual Ab. Discard 50 μL from the solution in column 10 after dilution so that each well of the 96-well plate has 50 μL of diluted Ab. Columns 11 and 12 are included as internal controls, as virus-only and mock-only, respectively. Each Ab is tested in triplicate.
4. In the BSL3, prepare ~100-200 PFU/well of SARS-CoV-2 in infectious media in the biosafety cabinet. From the virus stock, calculate ~1.0-2.0 x 10^4^ PFU and mix in 5 mL (for one 96-well plate) of infection media. Note: The amount of virus per well can be further optimized based on the virus titer in the stock.
5. Add 50 μL of SARS-CoV-2 to columns 1 to 11 of the Ab plate and incubate the mixture for 1 h at 37°C. Tap the plate gently to mix the sample with the virus.
6. After incubation, remove cell maintenance media from the Vero E6 96-well plate. Transfer 50 μL of the Ab sample-virus mixture from the 96-well plate to the corresponding Vero E6 in 96-well plate using a multi-channel pipette. Incubate for 1 h at 37°C in a 5% CO_2_ incubator to allow virus adsorption.
7. After 1 h, remove the virus inoculum and overlay with post-infection media containing 1% Avicel. Incubate the infected Vero E6 cells for 24 h at 37°C in the 5% CO_2_ incubator.
8. At 24 h post-infection (h p.i.), remove infectious media and fix/inactivate the plate in 10% formalin solution for 24 h at 4°C.

### Post-treatment PRMNT assay

1. Vero E6 cells are maintained as described in the previous pre-treatment protocol.
2. Seed ~1 x 10^4^ Vero E6 cells/well the day before virus infection in 96-well plates.
3. On the day of infection, check the confluency of the Vero E6 cells under a light microscope. The optimal cell confluency is between 85-95%.
4. Prepare a 2-fold serial dilution of the Ab or Ab containing sample in an empty, sterile 96-well cell culture plate using a post-infection media as described in step 4 before. Add 50 μL of 2% Avicel in post-infection media to each well containing the diluted Ab or media-only and no-virus control wells (columns 11 and 12, respectively) to give a final concentration of 1% Avicel in each well.
5. In the BSL3, prepare ~100-200 PFU/well of SARS-CoV-2 in infection media in the biosafety cabinet. From the virus stock, calculate 1.0-2.0 x 10^4^ PFU and mix in 5 mL (for one 96-well plate) of infection media.
6. Add 50 μL of virus inoculum to each well of the cell-cultured Vero E6 96-well plate from column 1 to 11, and incubate for 1 h at 37°C in the CO_2_ incubator for virus adsorption.
7. After 1 h adsorption, remove the virus inoculum and replace with 100 μL of post-infection media containing the serially-diluted Ab with 1% Avicel.
8. Incubate the cells for 24 h at 37°C in the CO_2_ incubator.
9. At 24 h p.i., remove media and fix/inactivate the plate in 10% formalin solution for 24 h at 4°C.

### PRMNT assay to identify compounds with antiviral activity against SARS-CoV-2

Several antiviral compounds are currently being evaluated for their effectiveness against SARS-CoV-2 (Buonaguro et al., 2020; Elfiky, 2020; Park et al., 2020b; Xu et al., 2020). Different antiviral compounds target different steps of the viral replication cycle which provides the basis for the use of a PRMNT assay for antiviral screening. Hydroxychloroquine, a potential antiviral compound has been suggested to inhibit SARS-CoV-2 by blocking fusion to the cell membrane, (Fox, 1993; Yao et al., 2020). Other compounds in clinical trials, such as remdesivir, target the viral RdRp which is involved in SARS-CoV-2 replication (Buonaguro et al., 2020). This protocol can be adopted to screen the inhibitory effect and concentration of anti-SARS-CoV-2 compounds irrespective of the viral replication step. Although this protocol describes the antiviral activity in post-infection conditions, an alternative assay where the antiviral drug is provided before viral infection will help to provide information on the pre-entry mechanism of antiviral activity of the compound. The antiviral activity is determined by assessing the effective concentration that inhibits virus replication in Vero E6 cells (96-well plates, ~4 x 10^4^ Vero E6 cells/well at confluency, triplicates) following viral incubation for 24 h. Note that the seeding density of Vero E6 in 96-well is ~1 x 10^4^ cells/well while cells at confluency is ~4 x 10^4^ cells/well. It is important to include as a positive control in this antiviral PRMNT assay a drug with a known antiviral activity against SARS-CoV-2, such as remdesivir (Elfiky, 2020). Unlike NAbs, some compounds could have cytotoxic effects. As a result, it becomes imperative to carry-out a cytotoxicity test (e.g. MTT assay) to evaluate for potential cytotoxicity of a given compound. Vero E6 cells for an MTT assay can be seeded on the same day as the cells for PRMNT assay so that the experiments can be conducted in parallel. Alternatively, our infrared staining technique can be used, concurrently, to evaluate cell viability in the same cells, in 96-well plates, used for PRMNT assay.

### A PRMNT assay to identify SARS-CoV-2 antivirals

1. Vero E6 cells are maintained as described in the previous protocol.
2. Seed ~1 x 10^4^ Vero E6 cells/well the day before the virus infection in 96-well plates, using triplicates for each of the drugs that will be tested.
3. On the day of infection, check the confluency of the Vero E6 cells under a light microscope. The optimal cell confluency is between ~85-95%.
4. Prepare a 2-fold serial dilution of the antiviral in an empty, sterile 96-well cell culture plates using post-infection media. Briefly, add 50 μL of infection media to columns 2 to 12, and add 100 μL of 2X of the desired starting concentration of each antiviral to column 1. The starting concentration of the antiviral will, generally, depend on the type of drug being evaluated. We normally use a starting concentration of 50 μM for antivirals. Transfer 50 μL of antiviral from column 1 to column 2, and mix ~ 10 times using a multi-channel pipette. Repeat this process from column 2 to column 10, changing the tips between dilutions to prevent transfer of residual antiviral. Discard 50 μL from the solution in column 10 after dilution so that each well of the 96-well plate has 50 μL of either diluted antiviral or no antiviral (for columns 11 and 12). Columns 11 and 12 are included as virus-only and mock-only control wells, respectively.
5. Add 50 μL of post-infection media containing 2% Avicel to each well in the 96-well plate.
6. Prepare ~100-200 PFU/well of virus in post-infection media in the biosafety cabinet. From the virus stock, calculate ~1.0-2.0 x 10^4^ PFU and mix in 5 mL (for one 96-well plate) of infection media.
7. Remove media from the cell cultured Vero E6 cells in the 96-well plate and add 50 μL of SARS-CoV-2 to columns 1 to 11. Incubate the plate for 1 h at 37°C in the CO_2_ incubator to allow viral adsorption.
8. After viral adsorption, remove viral inoculum from the 96-well plate. Transfer 100 μL of the serially diluted sample containing 1% Avicel to the corresponding wells in the cell-cultured Vero E6 96-well plate using a multi-channel pipette.
9. Incubate infected Vero E6 cells in the 96-well plate for 24 h at 37°C in the CO_2_ incubator.
10. At 24 h p.i., remove media and wash the cell gently to remove Avicel. Fix/inactivate the plates for 24 h at 4°C with 10% formalin solution.

### Development of the PRMNT assays

For development of the PRMNT assays described above, we have optimized two different protocols with comparatively similar results: the peroxidase and the infrared staining techniques. The infrared staining technique holds the advantage of its capability to measure cell viability in addition to measuring antiviral activity. In the case of the peroxidase staining, a separate assay is needed to assess cell toxicity of NAbs or antiviral compounds. Both assays rely on the use of the SARS-CoV-1 cross-reactive NP mAb, 1C7 (Amanat et al., 2020) but any other mAb targeting a SARS-CoV-2 viral antigen can be used in the peroxidase and the infrared staining described below.

### Peroxidase staining

1. Once the plates are in the BSL2 after inactivation, remove residual formalin solution by gentle wash with double distilled water (DDW) before staining. Be careful not to touch or dislodge the cells in the 96-well plate with pipette tips.
2. Gently wash the cells three times with 100 μL/well of PBS.
3. Permeabilize the cells with 100 μL/well of 0.5% Triton X-100 dissolved in PBS, and incubate at room temperature (RT) for 15 min in the biosafety cabinet. Note: If you use an Ab against a viral surface protein (e.g. S), you can skip this permeabilization step.
4. Wash the cells with 100 μL/well of PBS, three times, and block with 100 μL/well of 2.5% BSA in PBS. Incubate cells at 37°C for 1 h.
5. Prepare primary Ab solution (anti-NP mAb, 1C7, 1 μg/mL) in 1% BSA, in PBS. Add 50 μL/well of primary Ab solution and incubate at 37°C for 1 h.
6. After incubation with the primary Ab, wash each well three times with 100 μL/well of PBS.
7. Prepare the biotinylated anti-mouse Ab (VECTASTAIN^®^ ABC-HRP Kit, Peroxidase (Mouse IgG); Vector Laboratory) following the manufacturer’s instructions. For one 96-well plate, add 75 μL of normal blocking serum stock and 25 μL of biotinylated secondary Ab stock to 5 mL of PBS. Add 50 μL/well of biotinylated Ab solution to each well, and incubate for 30 min at 37°C.
8. Next, wash each well three times with 100 μL/well of PBS to remove biotinylated Ab solution thoroughly. Prepare VECTASTAIN ABC Reagent by following manufacturer’s instructions (VECTASTAIN^®^ ABC-HRP Kit, Peroxidase (Mouse IgG); Vector Laboratory). For one 96-well plate, add 50 μL each of Reagent A (Avidin, ABC) and Reagent B (Biotinylated HRP, ABC) to 5 mL of PBS. Then, add 50 μL/well of VECTASTAIN ABC Reagent and incubate for 30 min at 37°C.
9. Wash cells three times with 100 μL/well of PBS. Remove PBS, and dry the plate by gently blotting on paper towels. Prepare developing solution by following manufacturer’s instructions (DAB Substrate Kit, Peroxidase (HRP), with Nickel; Vector Laboratory).
10. Add 50 μL of developing solution to each well and wait until visible plaques are observed.
11. Stop the reaction by removing the developing solution, and wash with PBS. It is important not to wait for too long once the plaques are discernible to prevent the entire cells from turning black.
12. Take images and measure the stained positive cells using a CTL ImmunoSpot plate reader and counting software (Cellular Technology Limited, Cleveland, OH, USA). The formula to calculate percent viral infection for each concentration is given as [(Average # of plaques from each treated wells – average # of plaques from “no virus” wells)/(average # of plaques from “virus only” wells - average # of plaques from “no virus” wells)] x 100. A non-linear regression curve fit analysis over the dilution curve can be performed using GraphPad Prism to calculate NT_50_ or effective concentration 50 (EC_50_) of the Ab or antiviral (**Figures 2** and **5, respectively**).

**Figure 1.**
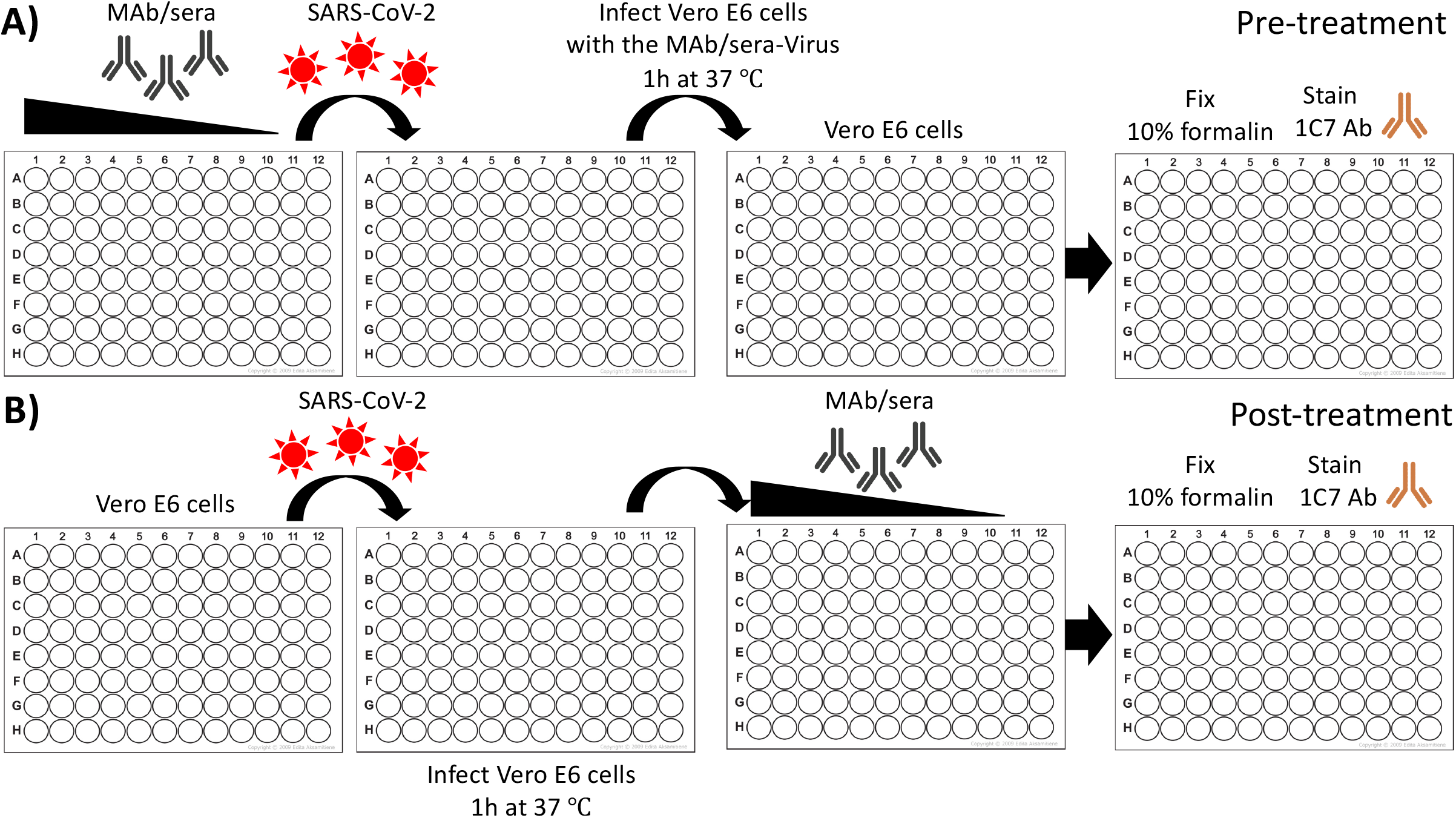
Schematic representation of the PRMNT assay to identify SARS-CoV-2 NAbs: Confluent monolayers of Vero E6 cells (96-well plate format, ~4 x 10^4^ cells/well, triplicates) are infected with ~100-200 PFU of SARS-CoV-2 pre-incubated **(A)** or post-treated **(B)** with a 2-fold serially diluted mAbs, pAbs, and/or serum samples. Cells in rows 11 and 12 are incubated as virus only and mock only infected cells, respectively, and are used as internal controls in the assay. After 1 h absorption of the virus-mAb mixture **(A)** or virus alone **(B),** infected cells are washed 3X with PBS and post-infection media containing 1% Avicel is added in all the wells alone **(A)** or with the 2-fold serially diluted mAbs and/or serum samples **(B).** At 24 h p.i., cells in the 96-well plate are fixed with 10% formalin solution. After 24 h, fixed cells are washed 3X with PBS and incubated with 1 μg/mL of a SARS-CoV-1 cross-reactive NP mAb (1C7, 1 μg/mL) at 37°C. After 1 h incubation with the primary Ab, cells are washed 3X with PBS and incubated with a secondary POD **(Fig. 2)** or IRDye 800CW goat anti-mouse IgG secondary Ab **(Fig. 3)** at 37°C. After 30 minutes incubation with the secondary Ab, cells are washed 3X with PBS and developed with the DAB substrate kit **(Fig. 2).** Cells stained with the IRDye 800CW goat anti-mouse IgG secondary Ab are simultaneously incubated with DRAQ5^™^ Fluorescent Probe Solution for nuclear staining **(Fig. 3).** Positive staining plaques in each of the wells of the 96-well plate is quantified using an ELISPOT plate reader **(Fig. 2)** or an Odyssey Sa Infrared Imaging System **(Fig. 3).** The neutralizing titer 50 (NT_50_) is calculated as the highest dilution of the mAb or sera that prevents 50% plaque formation in infected cells, determined by a sigmoidal dose response curve **(Figs. 2 and 3).**

**Figure 2.**
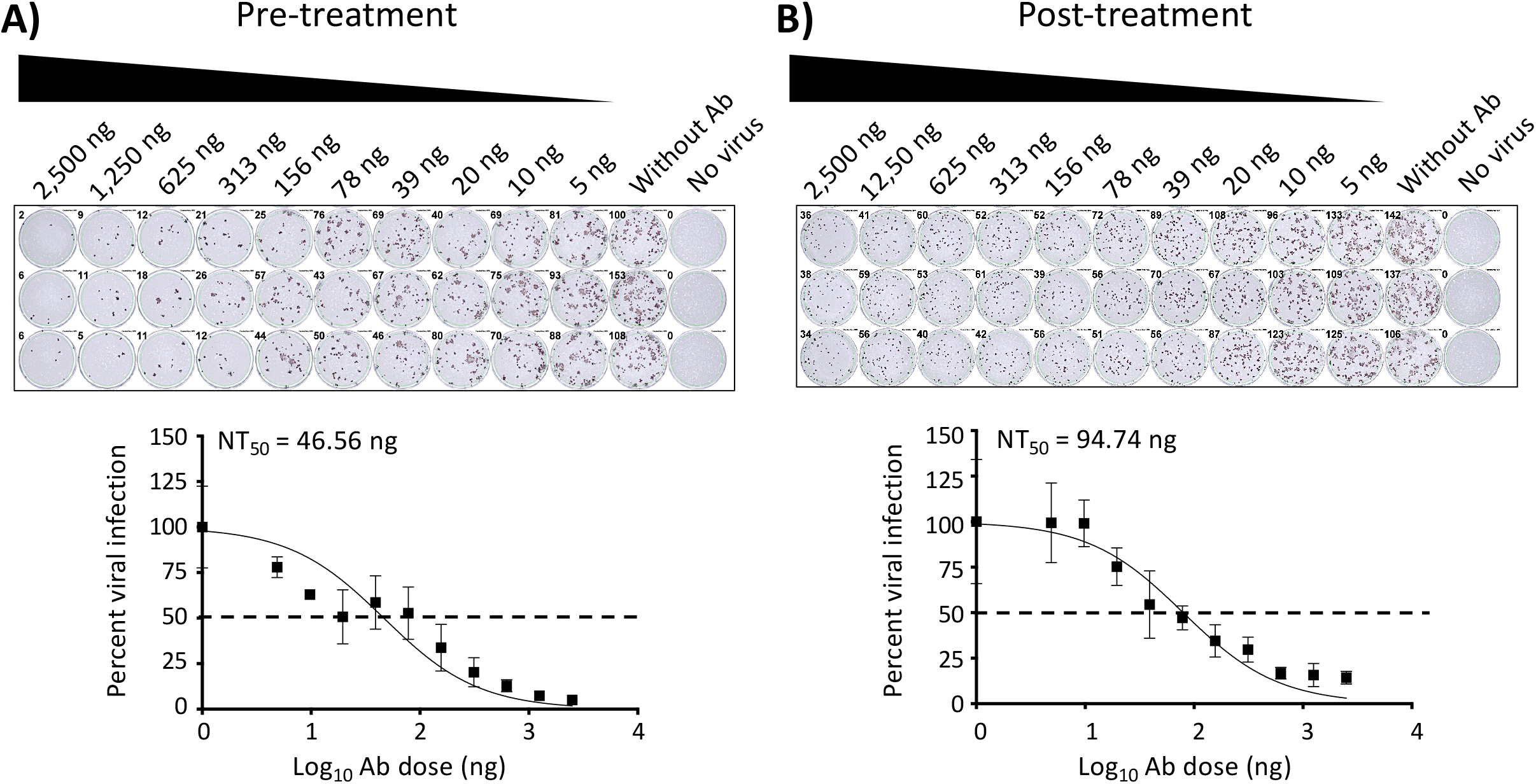
PRMNT assay to identify SARS-CoV-2 NAbs, peroxidase staining. A) Pre-treatment: 2-fold serially diluted mAbs, pAbs, or serum samples are pre-incubated with ~100-200 PFU of SARS-CoV-2 for 1 h at RT. After 1 h pre-incubation, confluent monolayers of Vero E6 cells (96-well plate format, ~4 x 10^4^ cells/well, triplicates) are infected with the mAb/pAb/serum-virus mixture for 1 h at RT. Cells in row 11 are incubated with virus only and cells in row are mock-infected and are used as internal controls in each of the plates. After 1 h of virus adsorption, cells are washed 3X with PBS and post-infection media containing 1% Avicel is added in all the wells. **B) Post-infection:** confluent monolayers of Vero E6 cells (96-well plate format, ~4 x 10^4^ cells/well, triplicates) are infected with ~100-200 PFU/well of SARS-CoV-2. After 1 h of virus adsorption, the infection media is replaced with post-infection media containing 1% Avicel and 2-fold serially diluted mAbs, pAbs or serum samples. Cells in row 11 are incubated with virus only and cells in row are mock-infected and are used as internal controls in each of the plates. **A and B)** At 24 h p.i., cells are fixed with 10% formalin solution. After 24 h fixation, cells are washed 3X with PBS and incubated with 1μg/mL of a SARS-CoV-1 cross-reactive NP mAb (1C7) at 37°C. After 1 h incubation with the primary mAb, cells are washed 3X with PBS and incubated with a secondary POD anti-mouse Ab (diluted according to the manufacturer’s instruction) at 37°C. After 30 minutes incubation with the secondary Ab, cells are washed 3X with PBS and developed with the DAB substrate kit. Positive staining plaques in each of the wells are quantified using am ELISPOT plate reader. The NT_50_ is calculated as the highest dilution of the mAb, pAb or sera that prevents 50% plaque formation in infected cells, determined by a sigmoidal dose response curve. Dotted line indicates 50% neutralization. Data were expressed as mean and SD from triplicate wells.

**Figure 3.**
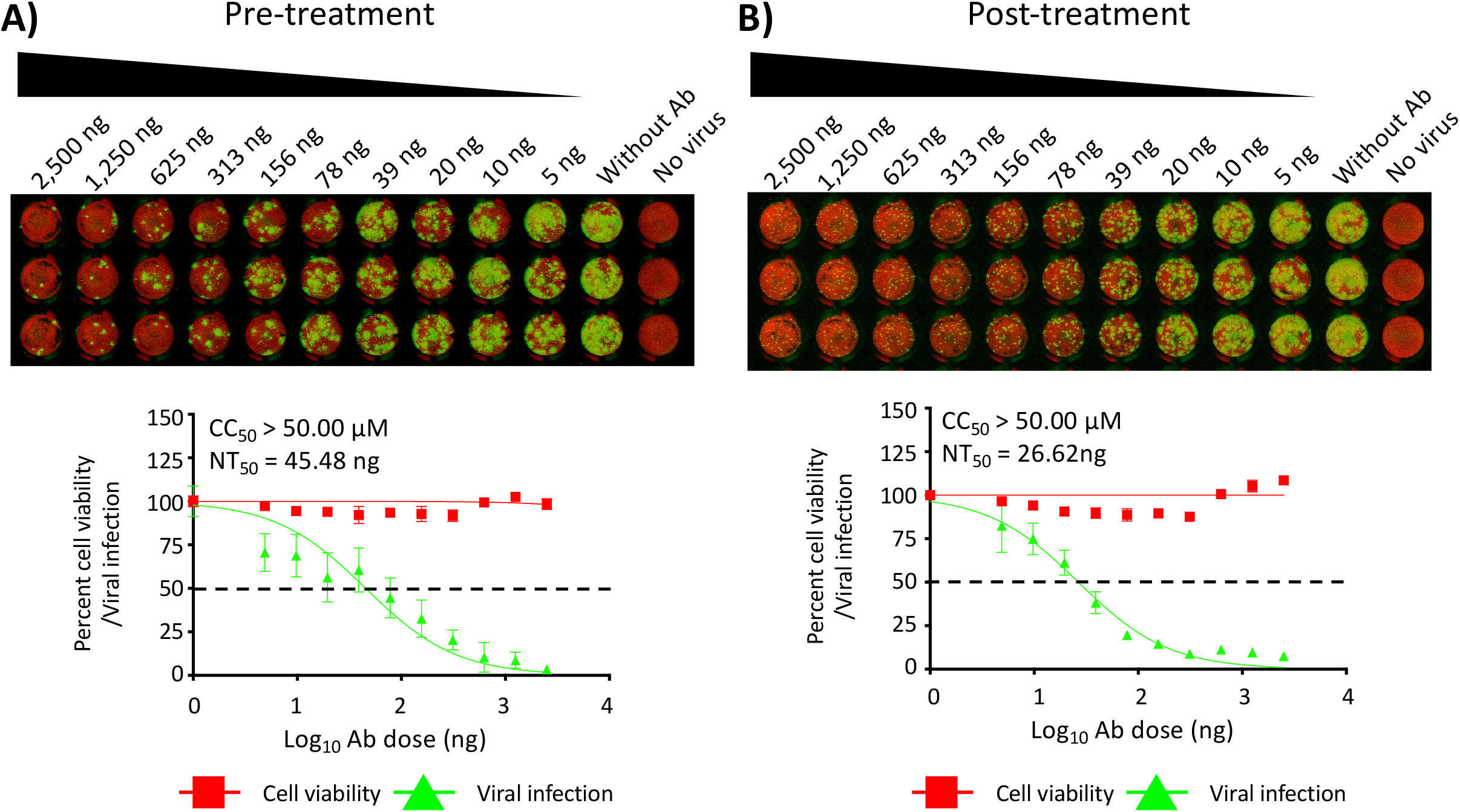
PRMNT assay to identify SARS-CoV-2 NAbs, fluorescent staining. **A-B)** Confluent monolayers of Vero E6 cells (96-well plate format, ~4 x 10^4^ cells/well, triplicates) are infected, fixed and staining with the primary NP 1C7 mAb as described in **Figure 2**. After incubation with the primary Ab, cells are incubated, for 1 h, with IRDye 800CW goat anti-mouse IgG secondary Ab, and DRAQ5^™^ Fluorescent Probe Solution for nuclear staining. After 1 h incubation with the secondary Ab and nuclear staining solution, cells are washed 3X with PBS and imaged using an Odyssey Sa infrared imaging system. The neutralizing titer 50 (NT_50_) is calculated as the highest dilution of the mAb, pAb, or sera that prevents 50% plaque formation in infected cells, determined by a sigmoidal dose response curve. Dotted line indicates 50% neutralization. Data were expressed as mean and SD from triplicate wells.

**Figure 4.**
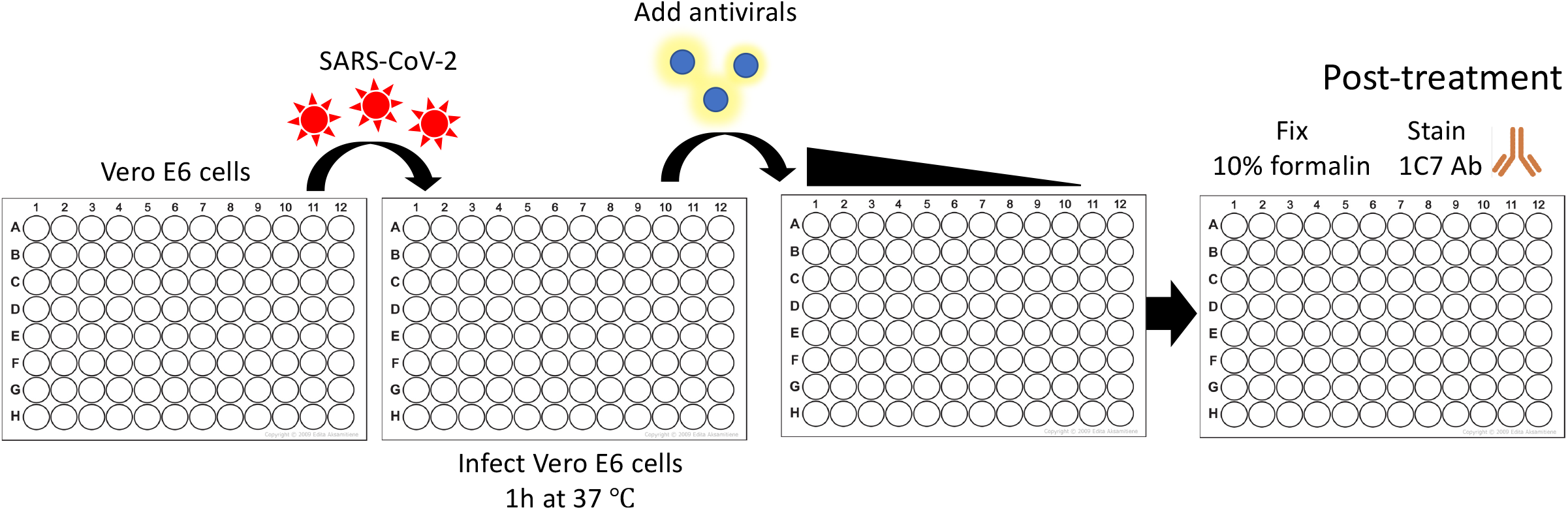
Schematic representation of the PRMNT assay to identify SARS-CoV-2 antivirals: Confluent monolayers of Vero E6 cells (96-well plate format, ~4 x 10^4^ cells/well, triplicates) are infected with ~100-200 PFU of SARS-CoV-2 for 1 h at 37°C. Cells in rows 11 and 12 are incubated with virus only and mock-infected, respectively, and are used as internal controls in each of the plates. After 1 h viral absorption, cells are incubated with post-infection media containing 2-fold serial dilutions of the antivirals containing 1% Avicel. At 24 h p.i., cells are fixed with 10% formalin solution. After 24 h fixation, cells are washed 3X with PBS and incubated with 1 μg/mL of the SARS-CoV-1 cross-reactive NP mAb (1C7) at 37°C. After 1 h incubation with the primary mAb, cells are washed 3X with PBS and incubated with a secondary POD (**Fig. 5**) or with IRDye 800CW goat anti-mouse IgG secondary Ab, and DRAQ5^™^ Fluorescent Probe Solution for nuclear staining (**Fig. 6**) at 37°C. After 30 minutes incubation with the secondary POD Ab, cells are washed 3X with PBS and developed with the DAB substrate kit (**Fig. 5**). Cells stained with the IRDye 800CW goat anti-mouse IgG secondary Ab are simultaneously incubated with DRAQ5^™^ Fluorescent Probe Solution for nuclear staining (**Fig. 6**). Positive stained cells in each of the wells of the 96-well plate are quantified using an ELISPOT plate reader (**Fig. 5**) or in the Odyssey Sa Infrared Imaging System (**Fig. 6**). The effective concentration 50 (EC_50_) is calculated as the highest dilution of the drug that prevents 50% plaque formation in infected cells, determined by a sigmoidal dose response curve (**Figs. 5 and 6**).

**Figure 5.**
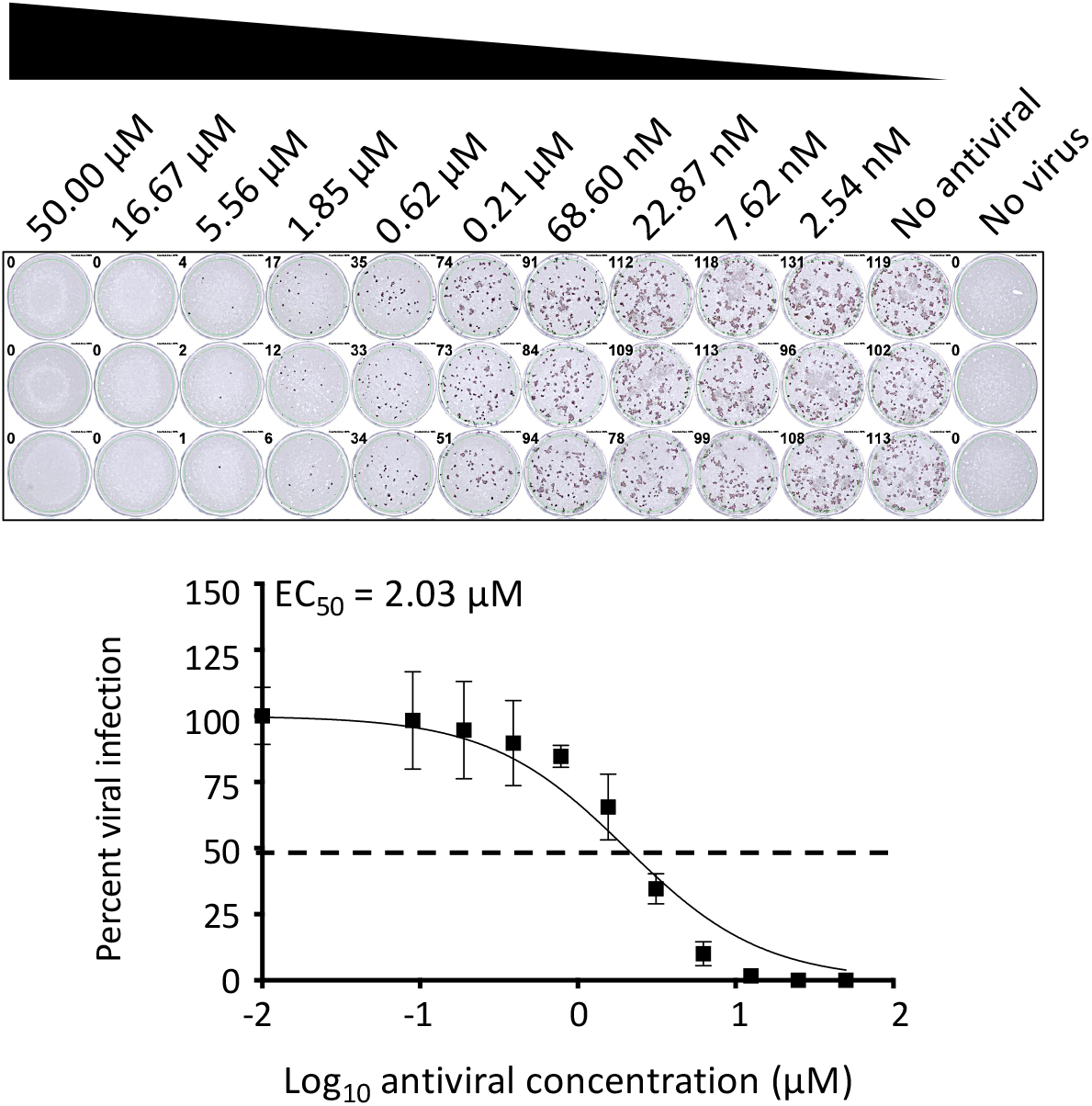
A PRMNT assay to identify SARS-CoV-2 antivirals, peroxidase staining: Confluent monolayers of Vero E6 cells (96-well plate format, ~4 x 10^4^ cells/well, triplicates) are infected with ~100-200 PFU/well of SARS-CoV-2. After 1 h of virus adsorption, media is replaced with fresh infection media containing 1% Avicel and 2-fold serially diluted drugs. Cells in row 11 are incubated with virus only and cells in row are mock-infected and are used as internal controls in each of the plates. At 24 h p.i., cells are fixed with 10% formalin solution. After 24 h fixation, cells are washed 3X with PBS and incubated with 1μg/mL of a SARS-CoV-1 cross-reactive NP mAb (1C7) at 37°C. After 1 h incubation with the primary mAb, cells are washed 3X with PBS and incubated with a secondary POD anti-mouse Ab (diluted according to the manufacturer’s instruction) at 37°C. After 30 minutes incubation with the secondary Ab, cells are washed 3X with PBS and developed with the DAB substrate kit. Positive staining plaques in each of the wells are quantified using an ELISPOT plate reader. The effective concentration 50 (EC_50_) is calculated as the highest dilution of the mAb, pAb or sera that prevents 50% plaque formation in infected cells, determined by a sigmoidal dose response curve. Dotted line indicates 50% neutralization. Data were expressed as mean and SD from triplicate wells.

### Infrared staining

1. Follow steps 1 to 6 as described for peroxidase staining.
2. After 1h of incubation with primary Ab (1C7, 1 μg/mL), wash cells three times with 100 μL/well of PBS. Next, prepare secondary Ab (IRDye 800CW goat anti-mouse IgG, 1:1,000 dilution) to stain virus-infected cells, and DRAQ5^™^ Fluorescent Probe Solution (1:4,000 dilution) to stain the nucleus using 1% BSA in PBS, and incubate for 1 h at RT.
3. Wash the cells three times with 100 μL/well of PBS to remove the secondary Ab and DRAQ5 thoroughly. Add 100 μL/well of PBS.
4. Obtain images and measure signal values of the stained positive cells with 700 nm (800CW, measuring viral infection) and 800 nm (DRAQ5, measuring cell viability) using Odyssey Sa Infrared Imaging System (LI-COR Biosciences, NE, USA). The formula to calculate percent viral infection (800 nm measurement) or cell viability (700 nm measurement) for each concentration is given as [(Average signal intensity from each treated wells – average signal intensity from “no virus” wells)/(average signal intensity from “virus only” wells - average signal intensity from “no virus” wells)] x 100. A non-linear regression curve fit analysis over the dilution curve can be performed using GraphPad Prism to calculate NT_50_ or EC_50_ of the Ab or antiviral (**Figures 3** and **6, respectively**).

**Figure 6.**
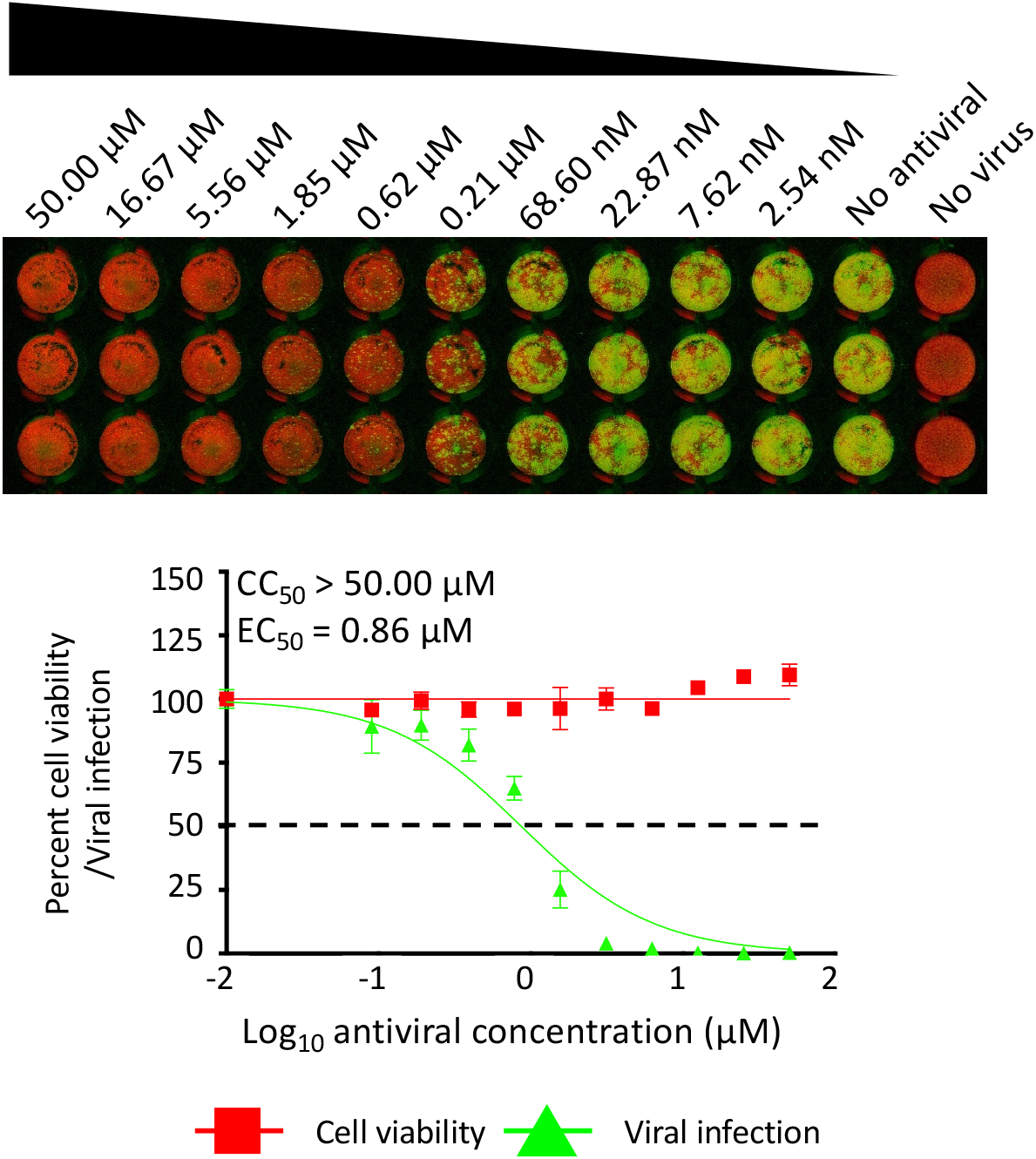
A PRMNT assay to identify SARS-CoV-2 antivirals, fluorescent staining: Confluent monolayers of Vero E6 cells (96-well plate format, ~4 x 10^4^ cells/well, triplicates) are infected, fixed and stained with the primary NP 1C7 mAb as described in **Figure 5**. After 1 h incubation with the primary NP 1C7 mAb, cells are washed 3X with PBS and incubated, at 37°C, with IRDye 800CW goat anti-mouse IgG secondary Ab (1:1000), and DRAQ5^™^ Fluorescent Probe Solution (1:4000) for nuclear staining. After 1 h incubation with the secondary Ab and nuclear stain, cells are washed 3X with PBS imaged using an Odyssey Sa Infrared Imaging System. The effective concentration 50 (EC_50_) is calculated as the highest dilution of the mAb, pAb or sera that prevents 50% plaque formation in infected cells, determined by a sigmoidal dose response curve. Dotted line indicates 50% neutralization. Data were expressed as mean and SD from triplicate wells.

## Discussion

As COVID-19 cases continue to increase across the globe, there is an urgent need to identify effective and safe prophylactics and/or therapeutics for the treatment of SARS-CoV-2 in infected patients. In this study, we described robust assays and staining techniques which can be employed to evaluate the neutralizing and/or antiviral activity of Abs and/or antivirals, respectively, against SARS-CoV-2 *in vitro*.

The PRMNT assay described here measures the neutralization effect of Abs or the antiviral activity of compounds against SARS-CoV-2 in similar manner to what we have previously described with other viruses, including, among others, influenza (Baker et al., 2015; Bauman et al., 2013; Martinez-Sobrido et al., 2010; Nogales et al., 2019; Nogales, Baker, and Martinez-Sobrido, 2015; Nogales et al., 2018; Nogales et al., 2016; Park et al., 2020a; Piepenbrink et al., 2019; Rodriguez, Nogales, and Martinez-Sobrido, 2017; Yang et al., 2016), mammarenaviruses (Brouillette et al., 2018; Ortiz-Riano et al., 2014; Robinson et al., 2016; Rodrigo, de la Torre, and Martinez-Sobrido, 2011), and Zika virus (Park et al., 2019). Although the assay described here was adapted to 96-well plate format, it can be adjusted to 384-well plate format for HTS of SARS-CoV-2 antivirals and/or NAbs. Many factors, including virus isolate, virus titer, cell condition or confluency, nature of samples, and incubation period can affect the outcome of this assay (Amanat et al., 2020). It is important to optimize these conditions so as to be able to generate accurate and reproducible data between different laboratories.

To restrict virus spread in infected cells, it is paramount to incubate the infected cells with post-infection media containing 1% Avicel. Incubation without Avicel will enhance a rapid cell-to-cell spread of virus resulting in higher NT_50_ or EC_50_. Although this assay works best in our hands using 1% Avicel, other reagents, such as 0.75% carboxymethylcellulose (CMC) can also be used (Oladunni et al., 2019).

Both the peroxidase and the infrared staining described here measures the level of SARS-CoV-2 infection based on the staining SARS-CoV-2 NP. However, the staining techniques can also be adapted for Abs that recognizes other SARS-CoV-2 proteins. While both staining techniques are comparable with regards to NT_50_ and EC_50_ values, the infrared staining offers the advantage of measuring virus replication and cell viability in a single assay, saving time and reagents needed to separately evaluate cytotoxicity.

## Acknowledgments

We want to thank BEI Resources for providing the SARS-CoV-2 USA-WA1/2020 isolate (NR-52281). We want to thank Maritza Quintero for her technical assistance with the infrared staining with the Odyssey Sa Infrared Imaging System. We would also like to thank members at our institutes for their efforts in keeping them fully operational during the COVID-19 pandemic and the IBC committee for reviewing our protocols in a time efficient manner. We would like to dedicate this manuscript to all COVID-19 victims and to all heroes battling this disease.

